# Machine learning guided rational design of a non-heme iron-based lysine dioxygenase improves its total turnover number

**DOI:** 10.1101/2024.06.04.597480

**Authors:** R. Hunter Wilson, Anoop R. Damodaran, Ambika Bhagi-Damodaran

## Abstract

Highly selective C-H functionalization remains an ongoing challenge in organic synthetic methodologies. Biocatalysts are robust tools for achieving these difficult chemical transformations. Biocatalyst engineering has often required directed evolution or structure-based rational design campaigns to improve their activities. In recent years, machine learning has been integrated into these workflows to improve the discovery of beneficial enzyme variants. In this work, we combine a structure-based machine-learning algorithm with classical molecular dynamics simulations to down select mutations for rational design of a non-heme iron-dependent lysine dioxygenase, LDO. This approach consistently resulted in functional LDO mutants and circumvents the need for extensive study of mutational activity before-hand. Our rationally designed single mutants purified with up to 2-fold higher yields than WT and displayed higher total turnover numbers (TTN). Combining five such single mutations into a pentamutant variant, LPNYI LDO, leads to a 40% improvement in the TTN (218±3) as compared to WT LDO (TTN = 160±2). Overall, this work offers a low-barrier approach for those seeking to synergize machine learning algorithms with pre-existing protein engineering strategies.

## Introduction

Selective and catalytic activation of aliphatic C-H bonds remains a long-standing challenge in synthetic chemistry.^[1–4]^ Biocatalysis has emerged as a potential solution as enzymes can perform C-H bond activation with high degrees of regio-, chemo-, and stereoselectivity.^[5–9]^ Late-stage, biocatalytic incorporation of desired functional groups streamline synthetic pathways and offer more sustainable solutions for challenging synthetic transformations. In particular, installation of hydroxyl groups into inert C-H bonds have been achieved with both heme-containing P450s and 2-oxoglutarate(2OG)-dependent non-heme iron metalloenzymes.^[10–15]^ Protein engineering efforts such as structure-based rational design or directed evolution have been implemented to improve yields and enzyme stability.^[16–21]^ Machine learning (ML) is being increasingly involved in protein engineering campaigns to decrease the overwhelming sampling of sequence space and produce desired reaction outcomes.^[22–26]^ Leveraging these new ML tools could enable more rapid discovery of beneficial mutations or starting templates for directed evolution. While ML tools can be integrated into engineering workflows, they often require extensive training datasets either from enzyme-specific reaction assays or non-structural genetic data.^[27–32]^ Tailoring of extensive datasets for each specific enzyme still requires burdensome front-work from researchers. Thus, synergizing ML technologies with existing rational design strategies offers a low-barrier solution for identifying potentially beneficial mutations in an engineering campaign.

Herein, we explore a ML-guided rational design strategy for the engineering of LDO, a recently characterized 2OG-dependent lysine dioxygenase (**Fig 1A**; referred to as ‘Hydrox’ in the previous work).^[33]^ LDO catalyzes the installation of a hydroxyl group into a chemically inert C-H bond on the C_4_ carbon of lysine. Hydroxylysine is a useful building block for high-value scaffolds in pharmaceutical and polymer industries; thus, engineering biocatalysts for its efficient production is desirable.^[34–39]^ By incorporating the neural network MutCompute^[40]^ algorithm in tandem with molecular dynamics (MD) simulations, we identify several mutations capable of increasing the total turnover number (TTN) of LDO. Overall, our ML-augmented rational design of LDO demonstrates a readily adoptable strategy for improving the catalytic performance of metalloenzyme biocatalysts.

**Fig. 1.**
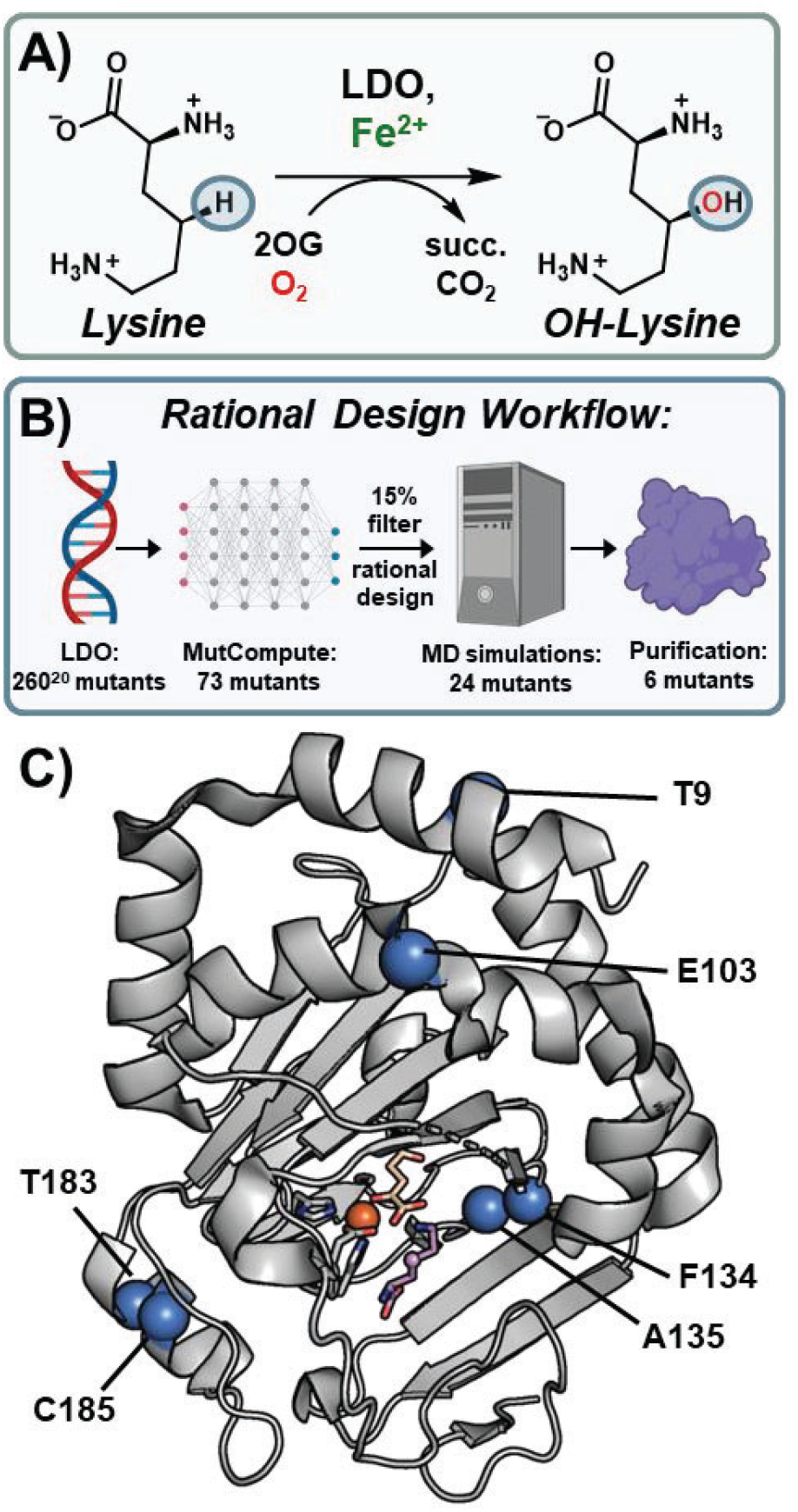
Machine learning-guided rational engineering of a non-heme iron 2OG-dependent dioxygenase. **A**) The non-heme iron enzyme, Lysine dioxygenase (LDO) catalyzes the hydroxylation of an aliphatic C-H bond at the C_4_ position of lysine. 2OG and molecular oxygen are co-substrates; succinate and CO_2_ are generated as byproducts. **B**) Rational design workflow overview starting from the total sequence space of LDO and ending with our six designed mutants. **C**) Crystal structure of LDO (PDB: 7JSD) showing ML-identified mutational hotspots as blue spheres. Iron displayed as an orange sphere, 2OG and Lysine are shown as beige and purple sticks, respectively.

## Results and Discussion

### Computational Design of LDO Variants

In the absence of extensive mutational reaction data with LDO, application of data-driven ML algorithms for designing variants with enhanced catalytic activity was not possible. Nevertheless, with the crystal structure of lysine-bound LDO in hand^[33]^, we could use structure-based ML algorithms to enhance its catalytic activity. We opted to use the MutCompute neural network, which was trained on >19000 sequence-balanced protein structures and can predict WT amino acid residues that are not optimized for their local environments. MutCompute has previously been shown to increase the turnovers and thermostability of several enzymes across a wide range of functions.^[26,41–43]^ Using the LDO crystal structure (PDB: 7JSD) as input, MutCompute identified 73 different mutational hotspots out of 260 total residues in the protein (**Fig. 1B**). Setting a cut-off filter of 15% WT probability, we down-selected 17 different mutational hotspots where the WT amino acid was disfavored (**Table S1**). We note that most of these amino acids are present on the surface of LDO. We inspected the crystal structure at the predicted mutational sites and developed rational design hypotheses for 24 potentially beneficial mutations (**Table S2**). MutCompute also provides suggested amino acid substitutions at each hotspot, which were included in our rational design strategy. We intentionally avoided mutational hotspots proximal (<12.5 Å) to the iron center, as it’s been demonstrated that enzyme activity in 2OG-depended non-heme iron enzymes can be greatly diminished by mutations proximal to the iron center.^[44–47]^ To verify potentially stabilizing interactions (due to increased H-bonding or hydrophobic packing interactions, etc.), we conducted MD simulations of our 24 rationally designed variants (**Table S2**). All mutational results were compared to simulations of WT LDO to discern whether new interactions were beneficial and did not diminish those present in the WT protein, to ultimately narrow down to six mutations for reactivity studies (**Fig. 1C**).

We began our MD analysis with the T9L mutation which was aimed to target the hydrophilic Thr9 residue placed in a hydrophobic pocket comprising of Leu10, Ileu6, Val99, and Leu49 (**Fig. 2A**). The T9L mutation was anticipated to increase hydrophobic packing and we observed this result by a lower temperature factor (B-factor) across MD simulations with this mutation when compared to the WT (**Fig. 2A inset**). Glu103 was the next target of our study which is located on a heavily strained alpha helix (**Fig S1**). We anticipated that mutating it to proline would help relax this strain and stabilize the protein. We did observe an average 0.5 Å lower root mean square deviation (RMSD) of the helix’s backbone atoms with E103P mutant as compared to WT LDO, suggesting this variant could relax the apparent strain. Phe134 was our next target residue which was placed in a charged/hydrophilic pocket comprising of Arg67, Asp137, Glu133, and Ser83 residues. We hypothesized that mutating this residue to a polar tyrosine would increase its stability within the charged/hydrophilic pocket while maintaining cation-π interactions with Arg67 (**Fig. 2B**). Indeed, we observe that the F134Y variant can form new H-bonds with water molecules present in the pocket (**Fig. 2B** inset). Next, substitution of Ala135 with a polar Asn was anticipated to form new H-bonds with residues on an adjacent loop (**Fig. S2**). Indeed, the MD simulations predicted new H-bonds with the side chains of Ser136 and Asp217. Additionally, mutation of this hydrophobic residue to a polar one increases H-bonds with water on the exterior of the protein. Residue Thr183 is located proximal to a local pocket surrounded by hydrophobic residues Ile180, Trp143, and Val168 (**Fig 2C**). Upon mutation to Ile, a lower pocket B-factor was observed in the MD simulations, suggesting that the T183I mutant would experience tighter hydrophobic packing. Finally, mutation of a solvent-exposed Cys185 to an isosteric, redox-inactive serine is predicted to increase H-bonding interactions with Asn181 and Glu182, located within the residing alpha helix (**Fig. S3**). Notably, all of these substitutions have a low predicted WT probability (out of the 20 proteogenic amino acids) as determined by MutCompute (**Table S1**). As all these residues are distal (>12.5 Å) from the LDO active site, they have the potential to form stabilizing interactions without sacrificing enzyme activity. Parsing through all residues distal from the active site would require time-consuming front-end labor. By incorporating the MutCompute neural network algorithm into this workflow, non-intuitive mutational hotspots were readily identified. Our MD simulations were instrumental in testing viable mutations created by rational design, as not all tested mutations resulted in stabilizing interactions (**Table S2**). For example, despite position Tyr18 having a low WT probability (1.9%), mutation of this residue to Phe (as suggested by MutCompute) resulted in the elimination of H-bonds with the backbone carbonyls of Met42/Arg43 as well as an increased B-factor of the local hydrophobic pocket (**Fig. S4**). Another design, N181W that also was suggested by MutCompute, resulted in a sharp increase in the B-factor of its adjacent hydrophobic pocket, weaking local hydrophobic interactions (**Fig S5**). Given these observed deleterious interactions, neither of these designs were expressed or purified. By incorporating newer ML algorithms with existing computational screening methods, we were able to propose six mutants for testing hydroxylation activity (**Fig. 1B**).

**Fig. 2.**
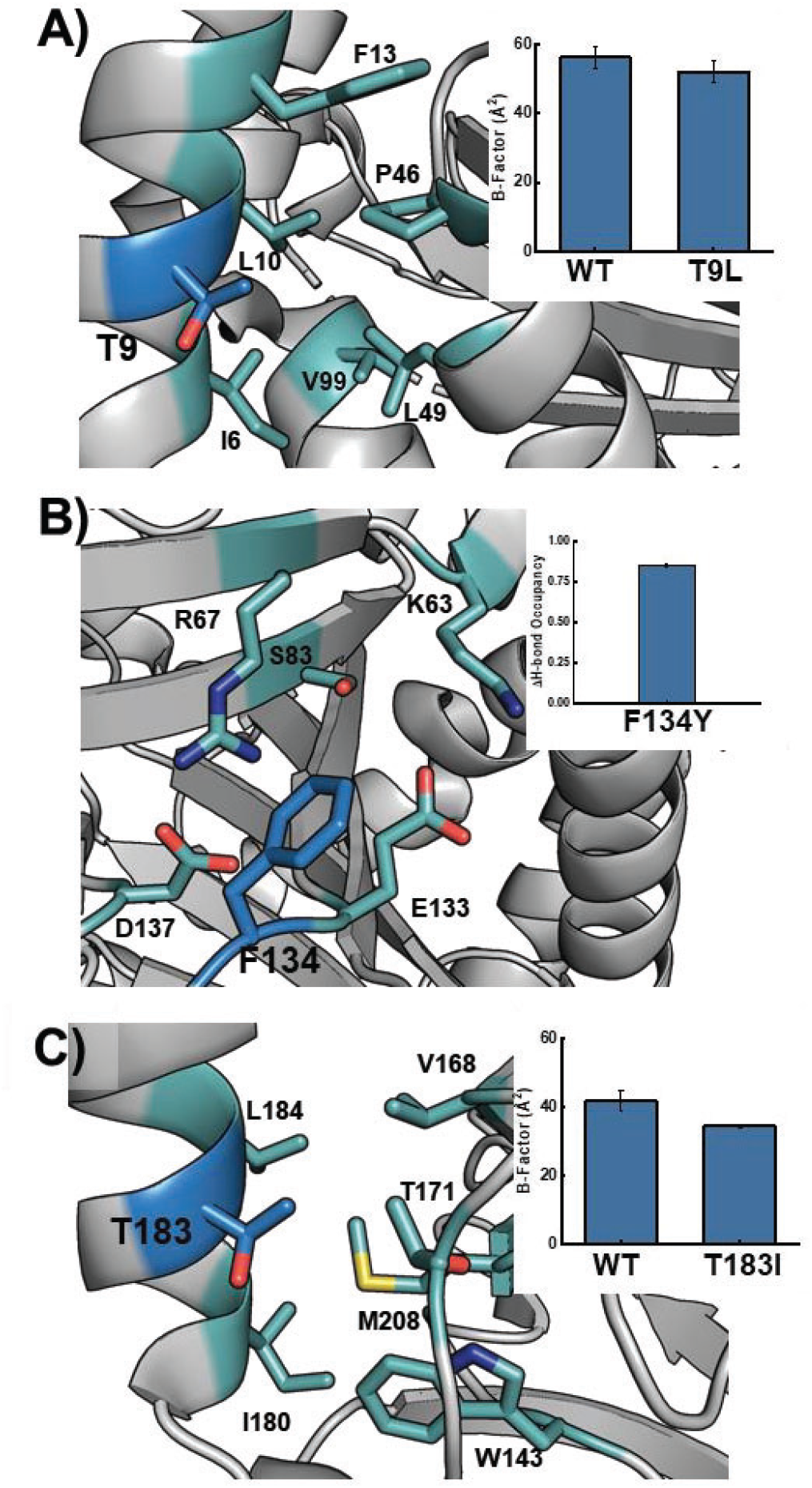
LDO variant design by Molecular Dynamics simulations. **A**) T9L variant (blue) can increase hydrophobic interactions by burying into an adjacent pocket (teal); inset: B-factor of pocket from MD simulations. **B**) Mutation of Phe134 (blue) to Tyr increases the stability within a highly polar pocket (cyan); inset: change in solvent H-bonding upon mutation. **C**) T183I mutant (blue) can bury into an adjacent hydrophobic pocket (cyan) to increase hydrophobic interactions; inset: B-factor of pocket from MD simulations.

### Purification of LDO Variants and Thermostability Analysis

WT LDO and all designed variants were expressed as His-Tagged constructs and purified by affinity chromatography. Interestingly, five of six variants were expressed in greater yields than WT, implying they exhibited increased solubility during expression conditions (**Fig S6**). The C185S mutant was isolated with approximately half the yield compared to WT, indicating that Cys185 may serve a structural role. Notably, Cys185 is the only cysteine residue in LDO, and it cannot form any intramolecular disulfide bonds. Disulfide bonds are also absent between monomeric chains in the crystal structure. Given that the other five mutations appeared to increase the protein yields, we also designed and purified a combinatorial mutant containing the other five mutations (T9L/E103P/F134Y/A135N/T183I), which we refer to as LPYNI LDO.

ML with MutCompute has been shown to enhance the thermostability of designed variants, resulting in increased melting points (T_m_) and turnovers.^[26,42,43]^ To assess the melting points of WT and designed LDO variants, we employed a thermal shift assay (TSA) to investigate how individual mutations impacted overall protein stability. In the apo forms of the protein, WT LDO had a T_m_ of 31.0°C with most of the other variants containing slightly lower T_m_ (within 2°C) (**Fig. S7**). Only A135N and C185S displayed melting points drastically lower than WT (-4.0°C), while T183I and the combinatorial LPYNI variant displayed elevated T_m_ values (+2.2 and +2.7°C, respectively). We note that since the variants were designed and simulated in a Fe/2OG-bound state, we would not know how our designs would behave in an unfolded state.

To validate our designs, we performed the TSA in the presence of metal and 2OG substrates to capture a state more similar to our simulations (**Fig. S8**). Interestingly, the T_m_ of the WT increases to 50.7°C, a +19.7°C shift relative to the apo form of the protein. This implies that metal and 2OG binding induce large-scale structural changes to form a stable, catalytically primed folded state. Lending credence to this, no crystal structure of a 2OG-dependent dioxygenase has been solved without an active site metal.^[48–54]^ A similar increase in T_m_ is observed across all variants, indicating that they can still bind cofactors necessary for catalysis. Relative to WT, most variants exhibit modest increases in T_m_ (+0.2-0.5°C for T9L, E103P, and F134Y) while T183I exhibits a T_m_ 3.5°C greater than WT. Analogous to the apo protein, A135N and C185S have lowered melting temperatures (-2.4°C and -4.2°C, respectively) relative to the ligand-bound WT. The combinatorial mutant, LPYNI displays a modest decrease in T_m_ of 0.9°C relative to WT, indicating that the stabilizing/destabilizing effects of individual mutations do not necessarily translate when combined with others.

We also investigated the effect of adding the substrate lysine to the protein complex (**Fig. 3A-C**). Upon binding the substrate, the melting point of the WT LDO protein complex increases to 54.5°C (+3.8°C relative to LDO without lysine). Once again, it appears that structural changes are occurring upon substrate binding which increase the stability of the overall complex. Recent structural studies in an analogous 2OG-dependent dioxygenase, HalD, demonstrated that upon substrate binding, a lid closes over the active site, securing the substrate’s orientation near the reactive iron center.^[55]^ Given our observed increase in melting temperatures upon lysine binding, a similar conformational change might be at present in LDO. Similar to WT, all mutants display an increase in T_m_ upon binding substrate lysine (**Figs. 3A-C, S8**). Most lysine bound mutants (T9L, E103P, and F134Y) display similar melting temperatures to the WT protein (within 0.5°C), while T183I displays an increased T_m_ (+3.0°C). While the quintuple LPYNI mutant displayed a slightly lower overall T_m_ than WT, the onset of melting was observed at temperatures higher than WT indicating higher stability under ambient conditions (**Fig. 3C**). Overall, these thermostability measurements reveal that these mutations do not necessarily enhance the T_m_ of the enzyme complex and show that all variants are capable of binding the substrates necessary for catalysis.

**Fig. 3.**
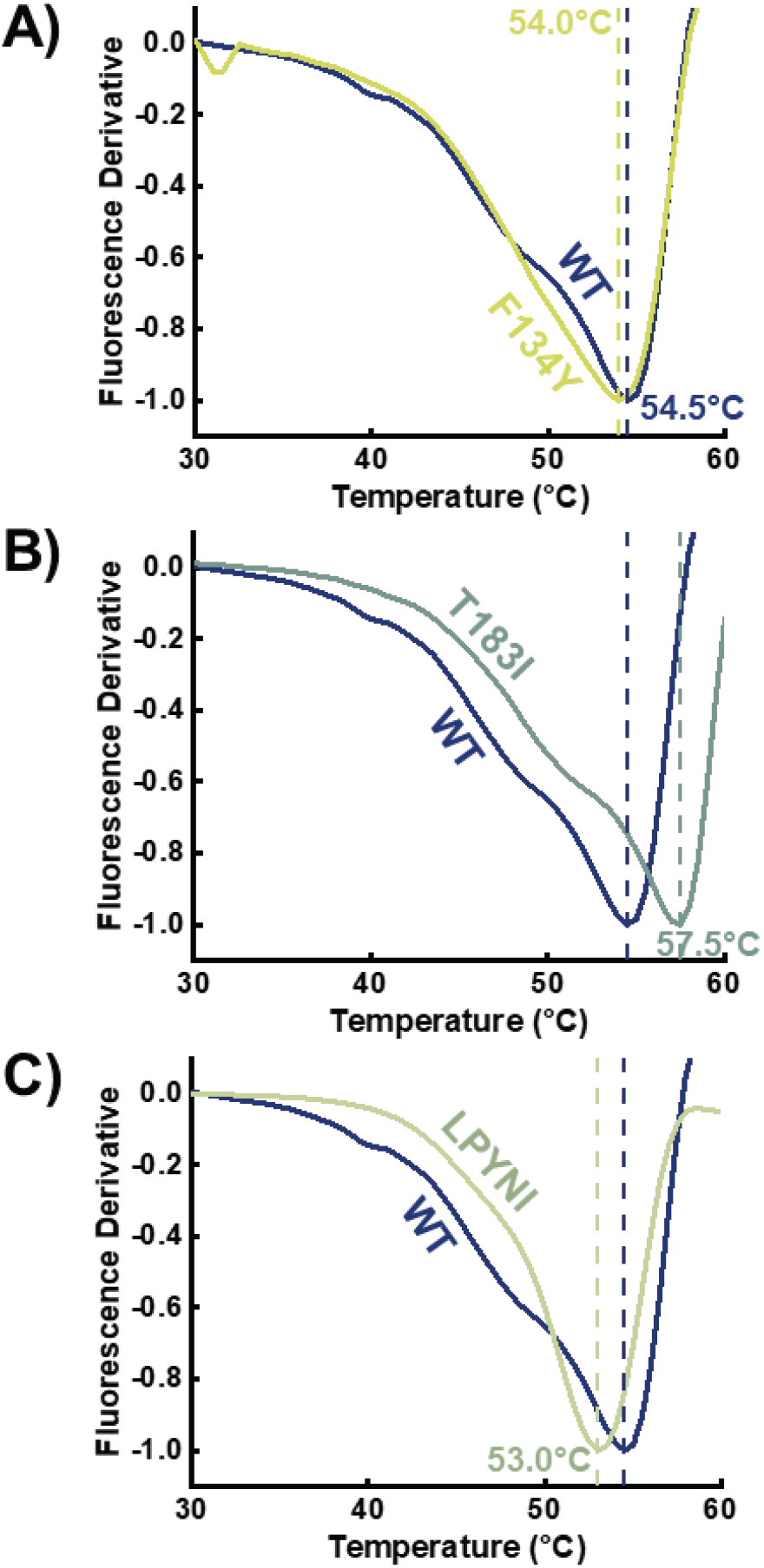
Melting point analysis for designed variants (**A** – F134Y, **B** – T183I, **C** – LPYNI) as compared to WT. All proteins were constituted with Mn^2+^, 2OG, and Lysine. Vertical lines are references for the WT and variant melting points.

### Catalytic Assessment of designed LDO Variants

While our TSA measurements indicate that the designed enzymes can bind their co-substrates, we wanted to assess their catalytic viability. While our TSA measurements indicate that the designed enzymes can bind their substrates, we wanted to assess their catalytic viability. Specifically, we wanted to assess their ability to hydroxylate lysine relative to the WT scaffold. To that end, we designed hydroxylation assays in which LDO variants were incubated with all necessary substrates (iron, 2OG, L-lysine) under aerobic conditions (**Fig. 1A**). After separation of protein from the reaction mixture, the lysine reactants and products were derivatized with a hydrophobic 6-aminoquinolyl-N-hydroxysuccinimidyl carbamate (AQC) tag. Derivatization with the AQC tag provides a chromophore for UV detection and enables retention of highly polar amino acids for reverse-phase HPLC analysis (**Fig. 4A**). In control reactions lacking LDO, a single peak for AQC-tagged lysine is present in the chromatogram (R_t_ = 4.8 min). The predominant peak at 2.2 min (Bis-AMQ) is a common byproduct of derivatization with AQC.^[56]^ In reactions incubated with WT LDO, a new peak appears at an earlier retention time (R_t_ = 4.3 min), indicative of the installation of a polar functional group into lysine. This implies the successful formation of OH-Lysine, which we validated with accurate mass measurements (**Fig S9**). The reaction product exhibited an m/z of 252.1055, with a mass error of 2.8 ppm, validating its identity. From the reaction, we determined a total turnover number (TTN) of 160±2 for WT Hydrox, which is in good agreement with the previously reported 136 as determined by mass spectrometric analysis.^[33]^

**Fig. 4.**
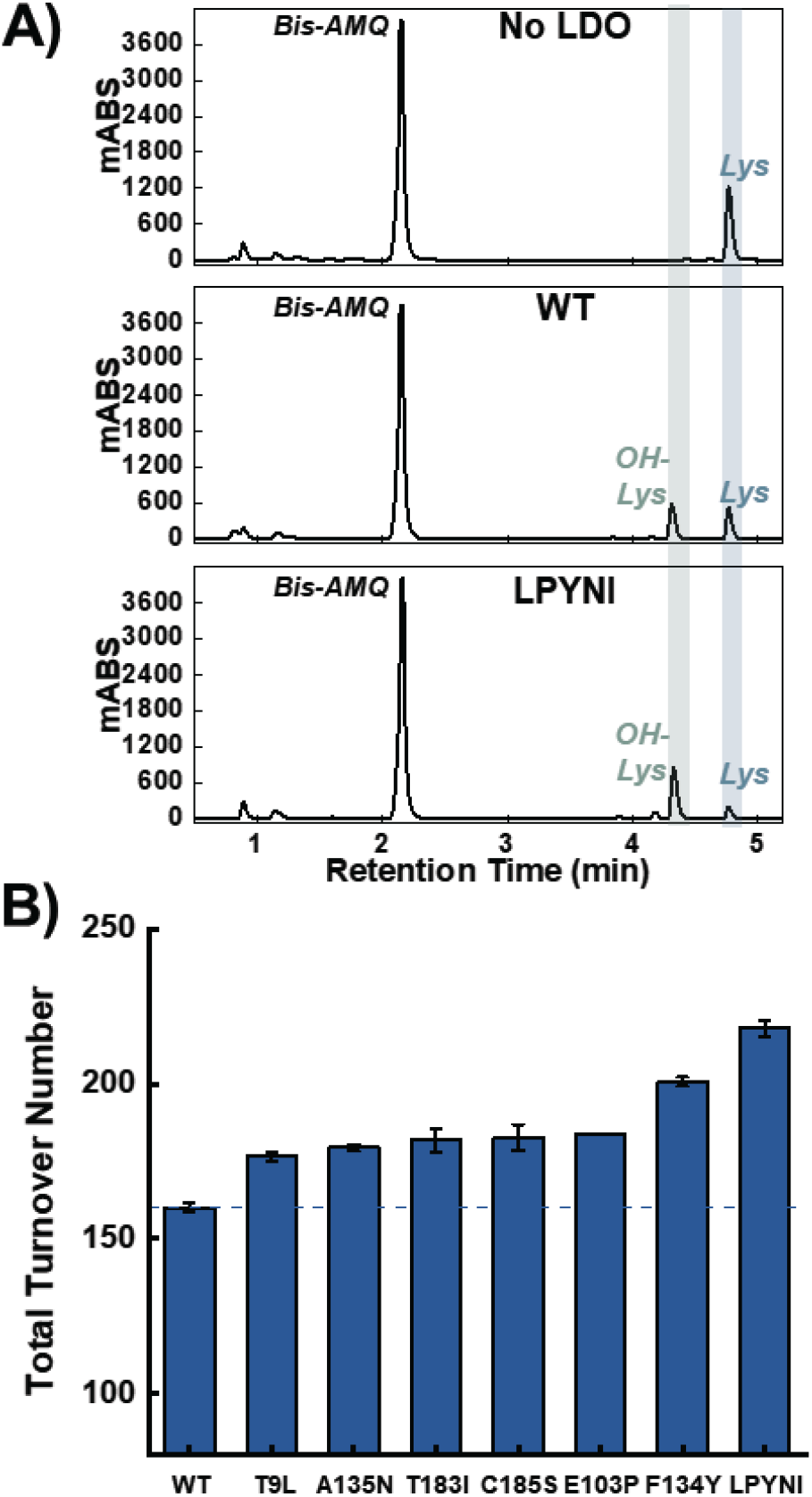
Hydroxylation assay analysis for LDO and designed variants. **A**) HPLC-UV detection of AQC-tagged lysine and hydroxylysine from a control reaction lacking LDO (top), a reaction with WT LDO (middle), and a reaction with LPYNI LDO (bottom). **B**) Total Turnover Number (TTN) for WT and LDO variants in the hydroxylation assay (n = 3). Horizontal line is a reference for the WT TTN. Error bars are the standard deviation from three independent reactions.

After establishing our assay with WT LDO, we tested how our rationally designed variants would affect the TTN upon reaction with LDO. HPLC chromatograms displayed lower amounts of reactant lysine and increased amounts of OH-lysine present in the reaction mixture relative to the WT reaction (**Fig. 4A**). Interestingly, all variants displayed a greater TTN than the WT enzyme (**Fig 4B**). Even mutations that decreased the T_m_ of LDO (A135N and C185S), resulted in increased turnovers over the WT LDO. While the individual mutants modestly increased TTN by 16-40 turnovers, we were intrigued as to whether the combinatorial LPYNI variant would perform better than the individual mutants. Indeed, LPYNI LDO displayed a TTN of 218±3, performing 1.4-fold better than WT. Despite this improvement in activity, we observe diminishing returns in TTN from the incorporation of individual mutants, analogous to results obtained from directed evolution campaigns. In a recent study, a 2OG-dependent hydroxylase was evolved to function on a non-native substrate after sampling thousands of variants for activity.^[19]^ While the increases in activity here can be perceived as marginal compared to gains from directed evolution, our engineering strategy still produces functional, improved enzymes with only 7 purified enzymes. Additionally, our rational design strategy circumvents the need for extensive front-end mutagenesis campaigns for generating training data for activity-specific ML algorithms. Overall, our rational design strategy results in biocatalysts capable of increased yields and more favorable TTN.

## Conclusion

We have developed a new ML-guided rational design strategy for enhancing the catalytic TTN of metalloenzymes. As a proof of concept, we targeted an iron/2OG-dependent hydroxylase which activates an inert C-H bond on the C_4_ carbon of lysine. By identifying mutational hotspots with MutCompute and simulating potential mutants with MD simulations, we were able to reduce the overwhelming sampling of sequence space and identify six potential mutations for purification (**Fig 1B**). The ML algorithm identified hotspots beyond the iron center enabling their discovery for rational design as well as preventing potentially deleterious mutations near the active site. Most mutants exhibited similar thermostability as the WT protein and all variants were able to bind the necessary substrates for catalysis. All designed variants displayed increased TTN, further validating our design methodology. While our present work focuses on engineering a 2OG-dependent hydroxylase, we anticipate that our rational design method could be applied for improving other members of this superfamily such as halogenases, cyclases, and desaturases. Beyond 2OG-dependent enzymes, this design method can be used towards engineering metalloenzymes containing more complicated cofactors such as porphyrins and iron-sulfur clusters.^[57,58]^ Overall, our rational design method presents an inexpensive and low-barrier method for enhancing the catalytic activity of functional biocatalysts.

## Supporting information

Supp Info ML-LDO

## Supporting Information

The authors have cited additional references within the Supporting Information.^[59–73]^

## Acknowledgements

RHW acknowledges the support of the National Institute of Health Chemical Biology Training Grant (T32GM132029). This work was supported by NSF CBET and CLP (Grant # 2046527). MutCompute prediction analysis was performed with an in-house program by Daniel Diaz, whom we acknowledge and thank for their assistance. Mass spectrometry analysis was performed at The University of Minnesota Department of Chemistry Mass Spectrometry Laboratory (MSL), supported by the Office of the Vice President of Research, College of Science and Engineering, and the Department of Chemistry at the University of Minnesota. The content of this work is the sole responsibility of the authors and does not represent endorsement by MSL personnel.

