## Supplementary material for "Machine learning guided rational design of a non-heme iron-based lysine dioxygenase improves its total turnover number": Supp Info ML-LDO

### Materials and Methods

#### General Experimental Details

The following chemicals were purchased from Millipore-Sigma (chemical grade in parentheses): Sodium Chloride (BioUltra), Imidazole (ACS Reagent), 2-mercaptoethanol (BME; BioUltra), Phenylmethanesulfonyl fluoride (Bioreagent), 2-oxoglutarate (99.7%), Iron(II) sulfate heptahydrate (Bioreagent), (+)-Sodium L-Ascorbate (Bioreagent), Sodium tetraborate (ACS reagent), Sodium Acetate (HPLC). Glycerol (molecular biology), L-lysine monohydrochloride, Ethylenediaminetetraacetic acid (EDTA; molecular biology), Calcium chloride (molecular biology), DNaseI, and Magnesium chloride (molecular biology) were all obtained from Fisher. (4-(2-hydroxyethyl)-1-piperazineethanesulfonic acid) (HEPES) buffer was procured from Dot Scientific. Dithiothreitol (DTT), RNaseI, and Isopropyl β-d-1-thiogalactopyranoside (IPTG) were obtained from GoldBio. All buffered solutions were prepared in MilliQ water obtained from a Barnstead GenPure water filtration system (Thermo Scientific) with a resistivity of at least 18.2 MΩ˙cm.

#### Plasmid design and site-directed mutagenesis

A codon-optimized DNA oligonucleotide containing the WT LDO gene was inserted into a pET-28a(+) vector using the custom gene services from Biomatik. Plasmid sequence/design was verified with Plasmidsaurus. The LDO construct was designed with an N-terminal $His_{10}$-Tag, a $(GSS)_2$ linker, and a PreScission Protease recognition site (in that order) before the start codon for the LDO sequence. The gene was codon-optimized for expression in E. coli. Site-directed mutagenesis for LDO variants was conducted with Phusion site-directed mutagenesis kit from Thermo Scientific. Primers are provided in Table S3. PCR experiments were performed using an Axygen MaxyGene II instrument as previously described.[59] PCR products were transformed into DH5α cells (Thermo Scientific) for plasmid expression. Single mutant DNA sequences were confirmed using Sanger sequencing with ACGT.

#### Protein Expression and Purification

Sequenced plasmids were transformed into BL21-DE3 Gold cells (Agilent) and grown on kanamycin plates. Single colonies from plates were used to inoculate an overnight primary culture (2XYT broth, 50 mL, [kanamycin] = 0.05 mg/mL). Secondary cultures (1 L 2XYT broth in 2.8 L flasks, [kanamycin] = 0.05 mg/mL, single drop of Antifoam agent (Sigma) included) were inoculated with 10 mL of primary culture and grown at 37°C till an $OD_{600}$ of 0.6-0.8 was reached. Cultures were cooled on ice bath for 15 minutes prior to induction with IPTG to a final concentration of 0.2 mM. Cultures were then shaken for 24 hours at 18°C prior to harvest by centrifugation. Cell pellets were flash frozen in liquid nitrogen and stored at -20°C till further use. This expression protocol was followed for both WT and mutant LDO variants.

Cell pellets were thawed in a water bath prior to resuspension in Buffer A (50 mM HEPES, 300 mM NaCl, 20 mM imidazole, pH = 7.5) supplemented with DNaseI, RNaseI, PMSF (1 mM), $MgCl_2$ (1 mM), and $CaCl_2$ (25 mM). The cell resuspension was sonicated for 20 minutes (30 s on, 15 s off cycle) and centrifuged to remove cellular debris (20000 RPM, 20 minutes, 4°C). Supernatant was filtered with 0.22 µm syringe filters and loaded onto a 5 mL HisTrapFF column (Cytiva, 2 mL/min binding rate) on an AKTA Start protein purification system (Cytiva). After loading, the column was washed with 12 CV of Buffer A supplemented with 10 mM BME. The column was further washed with 28% Buffer B (50 mM HEPES, 100 mM NaCl, 350 mM imidazole, pH = 7.5) for 10 CV. Recombinant protein was eluted with 10 CV of 100% Buffer B. Pooled elution fractions were transferred to washed dialysis membrane (6 kDa MWCO, Fisher) and PreScission Protease was added in a 1:50 protease:protein ratio. Protein was dialyzed overnight at 4°C in dialysis buffer (20 mM HEPES, 100 mM NaCl, 1 mM DTT, 1 mM EDTA, pH = 7.5). Protein was further dialyzed (twice) against dialysis buffer lacking EDTA for at least 2 hours to remove the chelator. Dialysate was filtered, spiked with imidazole to 20 mM, and reloaded onto the HisTrapFF column. His-Tag cleaved flow-through was collected. SDS-PAGE was performed to ensure full cleavage and >95% purity. The protein was concentrated in 10 kDa MWCO centrifugal filter units and glycerol was added to 5% volume. Protein concentrations were determined by the absorbance at 280 nm using extinction coefficients calculated from ProtParam. The protein was aliquoted, flash frozen in liquid nitrogen, and stored at -80°C till further use.

**Thermal Shift Assay**

LDO protein (15 µM) was incubated with or without $MnCl_2$ (50 mM), 2OG (100 mM), and L-lysine hydrochloride (1 M) in 50 mM Tris HCl buffer, pH = 7.5. Sypro Orange dye (Thermo Scientific) was present in a 3x final concentration (prepared from DMSO stock diluted in Tris HCl buffer pH = 7.5). Each sample well contained a final volume of 50 µL. Assays were conducted on a MyiQ2 rt-PCR detection system (Bio-Rad). The temperature was increased linearly by 0.5°C every 30 seconds. After incubation for 1 min, the fluorescence (standard FAM excitation/emission) was recorded. Melting temperatures ($T_m$) were found by finding the global minimum value in the negative first derivative of the raw fluorescence data. Error bars were generated from the standard deviation in $T_m$ from three biological replicates.

**Lysine Hydroxylation Assay**

In reaction volumes of 100 µL, L-Lysine hydrochloride (3 mM), ferrous sulfate heptahydrate (1 mM), sodium L-ascorbate (5 mM), 2OG (5 mM) in 20 mM HEPES buffer (pH = 7.5) were incubated in an Eppendorf Thermomixer at 20°C. Reactions were initiated by the addition of LDO protein to a 10 µM final concentration. Reactions were spun at 400 RPM for four hours. Holes were punched in the top of the eppendorf tubes containing the reaction mixtures to enable facile gas diffusion. For timepoint experiments, 100 µL reaction aliquots were quenched by mixing with 10 µL of 100 mM EDTA. To isolate small molecules from protein, the reactions were concentrated in 0.5 mL 10 kDa MWCO centrifugal filter units and the flow-through was stored at -20°C till derivatization.

### Amino acid derivatization and HPLC analysis

Amino acid product flow through (7 µL) was mixed into 12.5 mM sodium tetraborate buffer (173 µL, pH = 10.5) in silanized, glass Waters vials. Derivatization was accomplished by adding 20 µL of 6-aminoquinolyl-n-hydroxysuccinimidyl carbamate (AQC, 3 mg/mL in acetonitrile; Cayman Chemical) and the solution was vortexed for 30 seconds. These reaction conditions ensure that both amine groups of lysine are AQC-derivatized.

HPLC analysis was performed on a Shimadzu Prominence-i LC-2030C 3D Plus system equipped with a InfinityLab Poroshell 120 CS-C18 column (3.0 mm x 150 mm x 2.7 µm; Agilent) and a InfinityLab Poroshell 120 CS-C18 guard column (3.0 mm x 5 mm x 2.7 µm; Agilent). Mobile phase A was 5 mM ammonium acetate (pH = 5.0) and mobile phase B was 60% Acetonitrile / 40% water. AQC-tagged analytes were eluted using the following method: 0-2 min, 20%B; 2-5.5 min; 75%B; 5.5-6 min, 100%B; 6-7 min, 100%B; 7-7.5 min, 100%B; 7.5-10 min, 20%B. AQC-tagged products were detected using the absorbance at 254 nm. Integrated peak areas were related back to concentration by a standard curve of L-lysine. After multiplying by the dilution factor, the concentration of products was divided by the LDO protein concentration to afford the total turnover number.

### Machine-Learning Guided Rational Design

The crystal structure for LDO (PDB: 7JSD) was used as input for the MutCompute neural network using a custom version of the neural network. Outputs from MutCompute provide the WT probability score for each position in the crystal structure. Positions with low WT probability were selected as hotspots (**Table S1**). If the hotspot was located within the protein active site or the amino acid was forming interactions with amino acids on a separate protein chain, then those positions were excluded from modelling. After inspection of the local protein environment surrounding the hotspot, several potential mutations were rationalized. MutCompute provides recommendations for mutations which were also considered. Mutant designs were screened by Molecular Dynamics simulations.

**Molecular Dynamics Simulations of Rationally Designed Mutants**

The starting structure of LDO was taken from the protein data bank (PDB: 7JSD; Chain C). Missing residues were added using the MODELLER module in UCSF Chimera.[60–64] In the tleap module, LDO was parameterized with the AMBER ff19SB force field, 2OG was treated with the generalized amber force field, and iron was treated as a ferrous species.[65–69] Force field parameters describing covalent bonds to iron were generated with the MCPB.py module.[70] L-Lysine was described using AMBER zwiterrionic amino acid parameters. The protein was solvated in a 10.0 Å OPC water box, and counterions $Na^+$ and $Cl^-$ were added to neutralize the system.[71] Mutations were performed by manually editing the input PDB files, deleting all atoms up to the beta carbon and renaming prior to input file generation with tleap. Proteins were minimized (first solvent, then protein and solvent), gently heated to 300 K, and density equilibrated for 2 ns. For WT and each mutant, three independent 100 ns production trajectories were developed. Trajectory analysis was performed with CPPTRAJ.[72] H-bond donor/acceptor distance cutoff were set to 3.2 Å. Hydrophobic interactions were monitored by measuring the B-factor of all amino acid side chains within a local pocket (keyword: molsurf). Error bars were calculated using the standard error over the three simulations (standard deviation divided by $\sqrt{3}$).[73] When calculating the change in H-bonding occupancy between WT and mutants, the error bars were calculated by propagating the error between WT and the mutant standard errors.

**Table S1**. MutCompute predicted hotspots for possible mutation. Probabilities for the WT amino acid are out of a maximum of 1 (ie. 100% probability).

| LDO AA position | WT Probability |
| --- | --- |
| T9 | 0.039 |
| Y18 | 0.019 |
| H41 | 0.017 |
| M42 | 0.120 |
| Q88 | 0.071 |
| D89 | 0.025 |
| D94 | 0.080 |
| E103 | 0.025 |
| F134 | 0.063 |
| A135 | 0.021 |
| T138 | 0.039 |
| N181 | 0.005 |
| T183 | 0.021 |
| C185 | 0.130 |
| V195 | 0.018 |
| S204 | 0.055 |
| D217 | 0.049 |

**Table S2**. LDO Mutants screened by Molecular Dynamics Simulations. Those demonstrating new or enhanced interactions were expressed and purified.

| Mutation | Expressed & Purified |
|---|---|
| T9L | Y |
| T9R | N |
| Y18F | N |
| H41E | N |
| H41S | N |
| M42E | N |
| Q88K | N |
| D89E | N |
| D94T | N |
| E103P | Y |
| F134Y | Y |
| A135N | Y |
| T138E | N |
| N181P | N |
| N181W | N |
| T183I | Y |
| C185S | Y |
| C185K | N |
| V195N | N |
| V195K | N |
| V195Q | N |
| V195R | N |
| D217P | N |

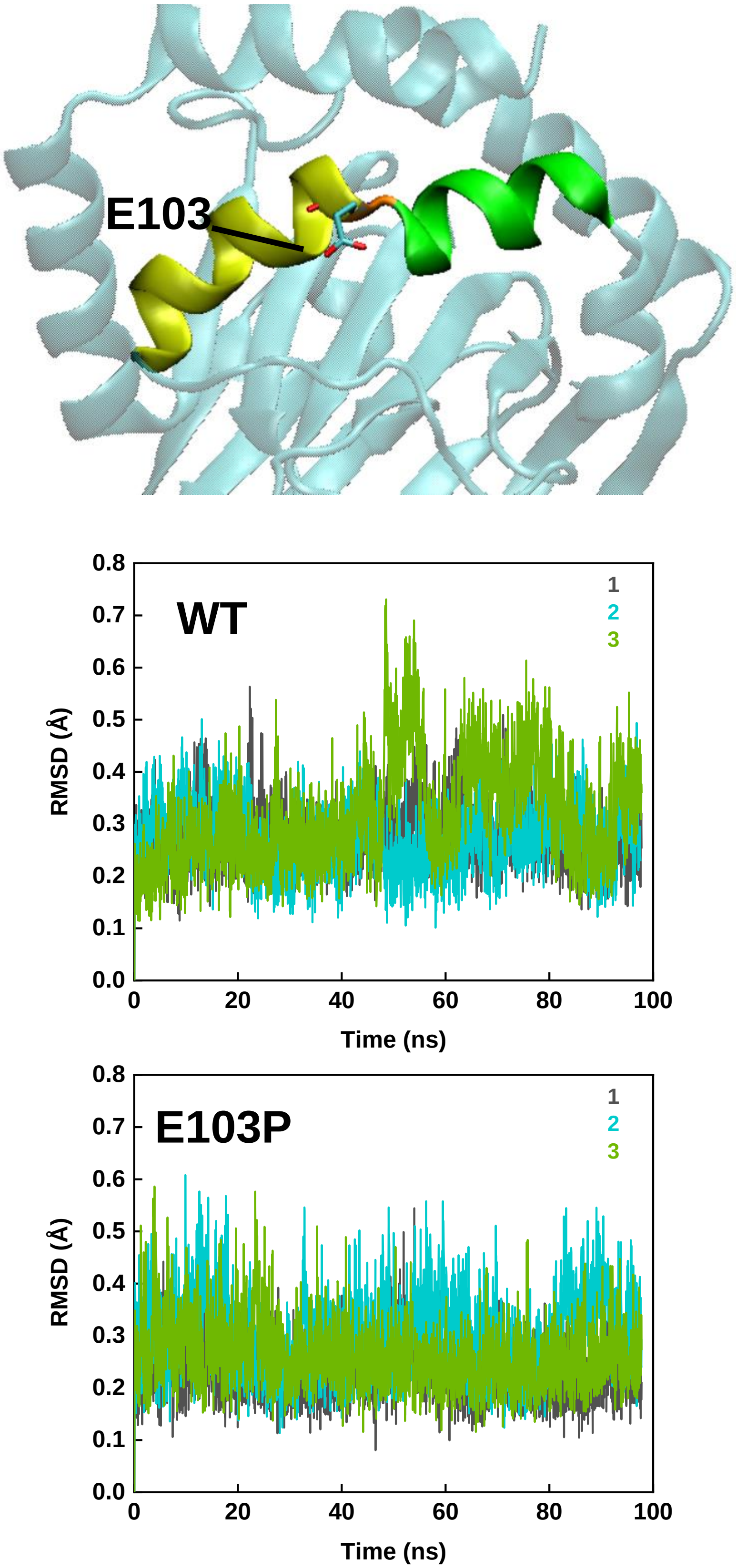


**Figure S1**. Designed E103P LDO variant. Glu103 is located in the middle of a highly strained alpha helix (either side highlighted in yellow/green; top panel). Monitoring the RMSD of the backbone atoms along the helix (residues 100-107), we can determine the overall stability of the helix. On average, the RMSD of the WT (middle panel) is 0.5 Å greater than E103P (bottom panel), indicating the mutation has stabilized the helix.

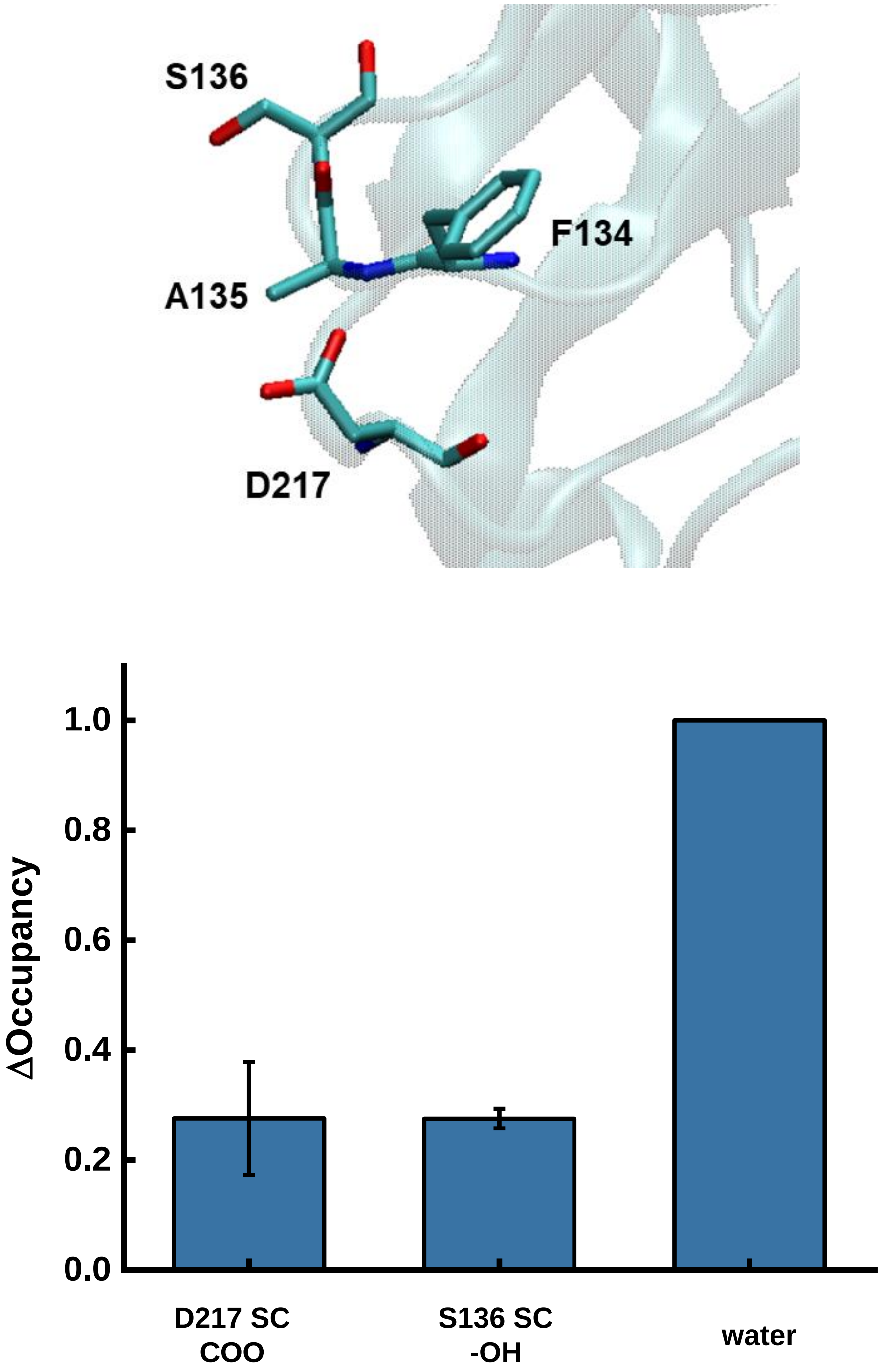


**Figure S2**. Designed A135N LDO variant. Ala135 is located on a loop proximal to polar and charged residues (top panel). Mutation to polar Asn enables H-bonding with neighboring residues and enhances solvent accessibility (bottom panel). Change in occupancy reflects the fractional change in a specific H-bond occurring upon mutation relative to WT. Error bars are the propagated standard error across three independent simulations SC = side chain.

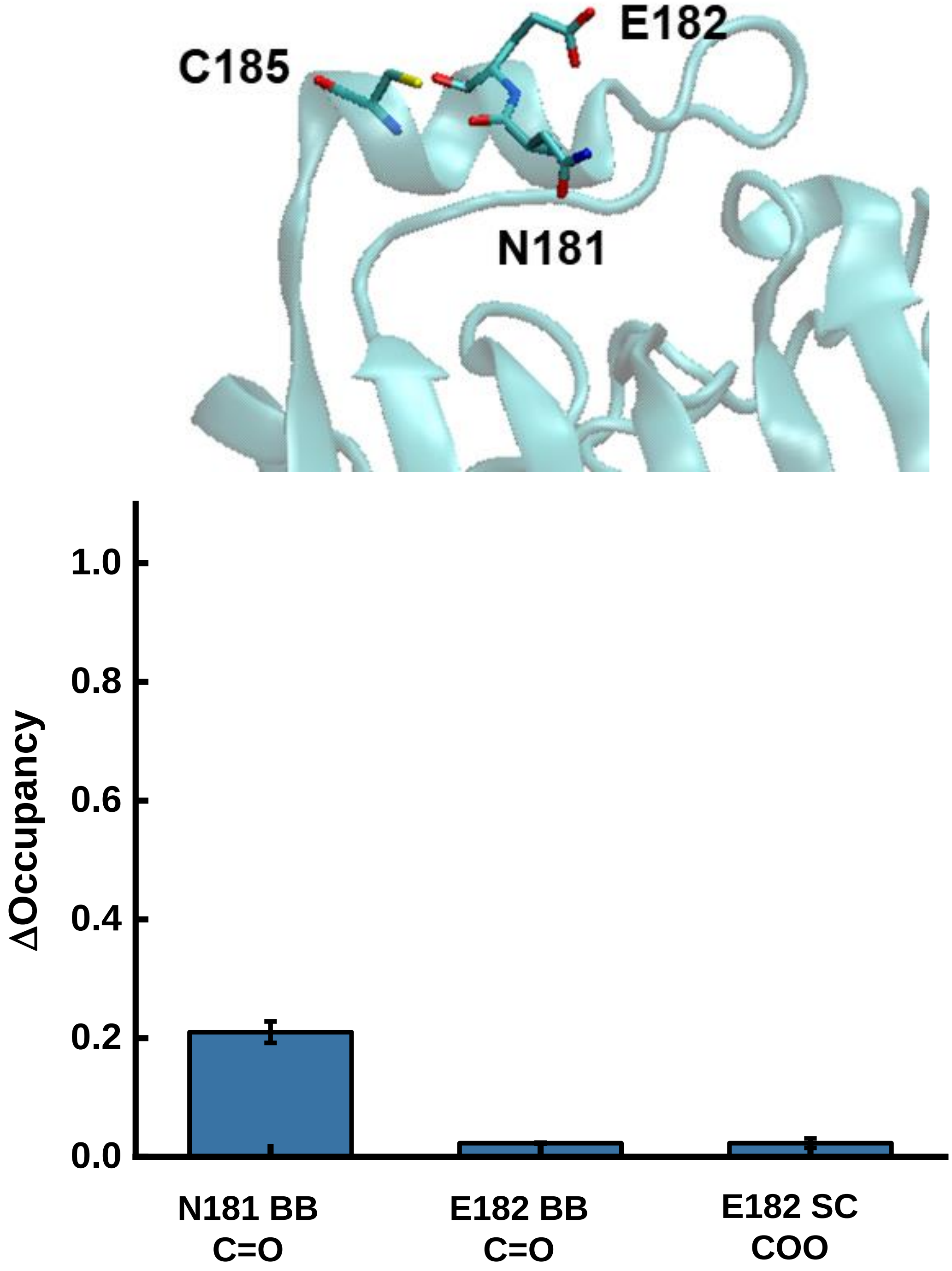


**Figure S3**. Designed C185S LDO variant. Cys185 is located on an alpha helix, solvent exposed, and a poor H-bond donor (top panel). Mutation to an isosteric, redox-inactive serine could enhance H-bonding and potentially decrease protein aggregation due to disulfide bond formation (bottom panel). Change in occupancy reflects the fractional change in a specific H-bond occurring upon mutation relative to WT. Error bars are the propagated standard error across three independent simulations. BB = backbone; SC = side chain

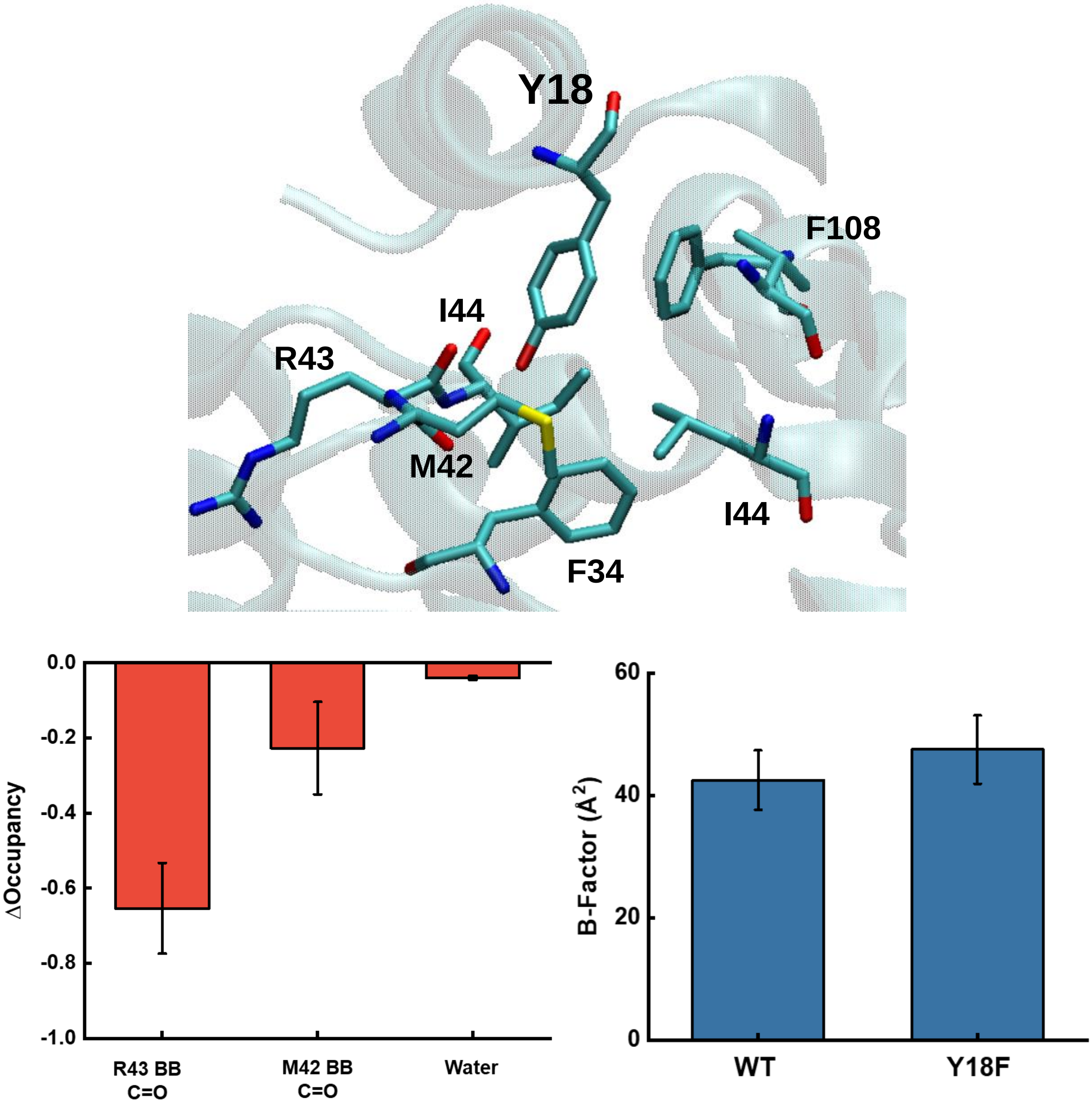


**Figure S4**. Designed Y18F LDO variant. Tyr18 is located on an alpha helix and located adjacent to a nearby hydrophobic pocket. Upon mutation to Phe, key H-bonds to Met42 and Arg43 were eliminated, and the pocket B-factor increases. Given these disruptions in the local environment, we did not move forward with this mutant. Change in occupancy reflects the fractional change in a specific H-bond occurring upon mutation relative to WT. Error bars are the propagated standard error across three independent simulations for H-bonding and the standard error for the B-factor data. BB = backbone

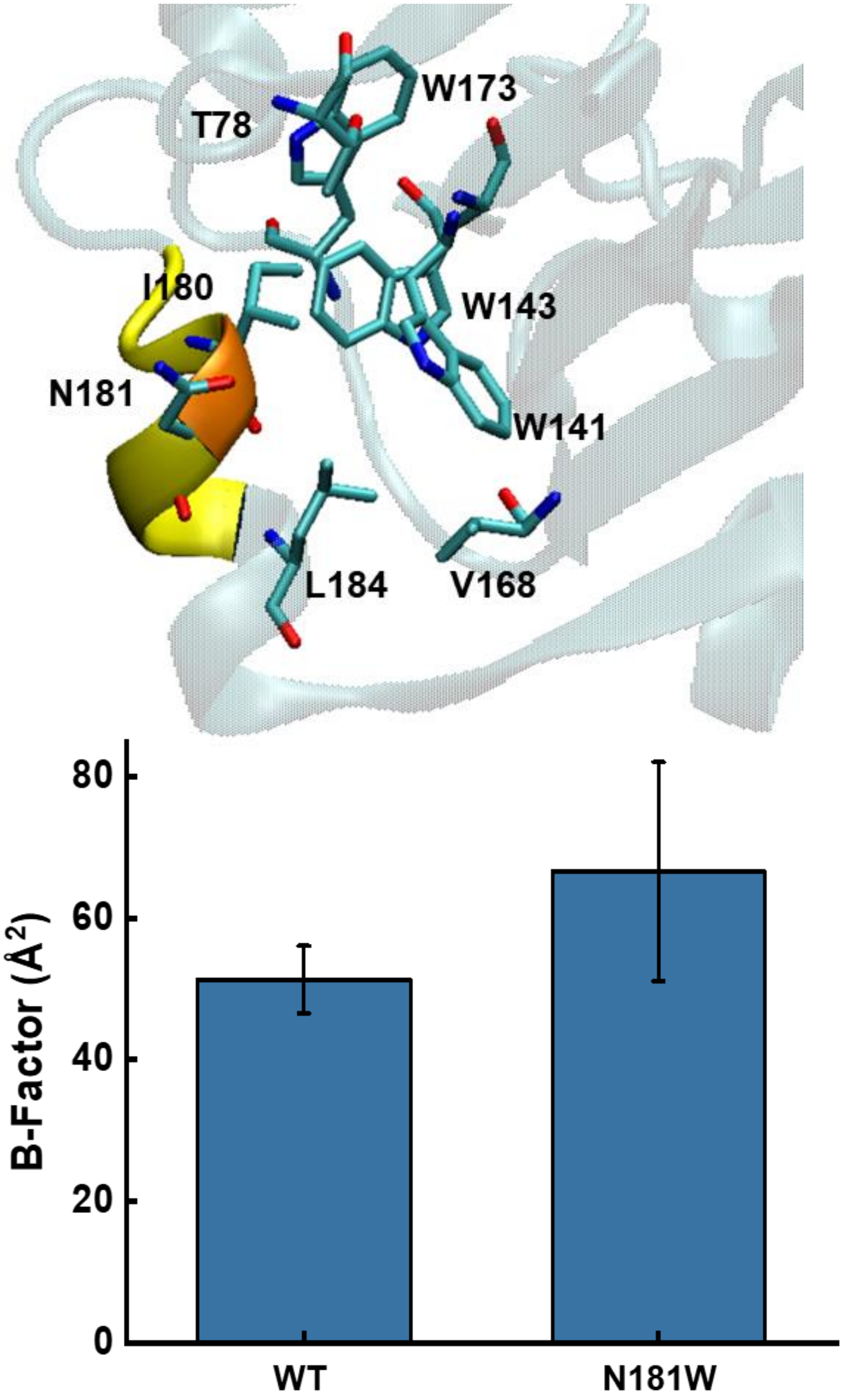


**Figure S5**. Designed N181W LDO mutant. Polar Asn181 resides on the surface of LDO near an adjacent hydrophobic pocket. Mutation to hydrophobic Trp results in an increase in B-factor suggesting poor hydrophobic packing. Given this decreased stability, we did not move forward with this mutant. Error bars are the standard error across the simulations.

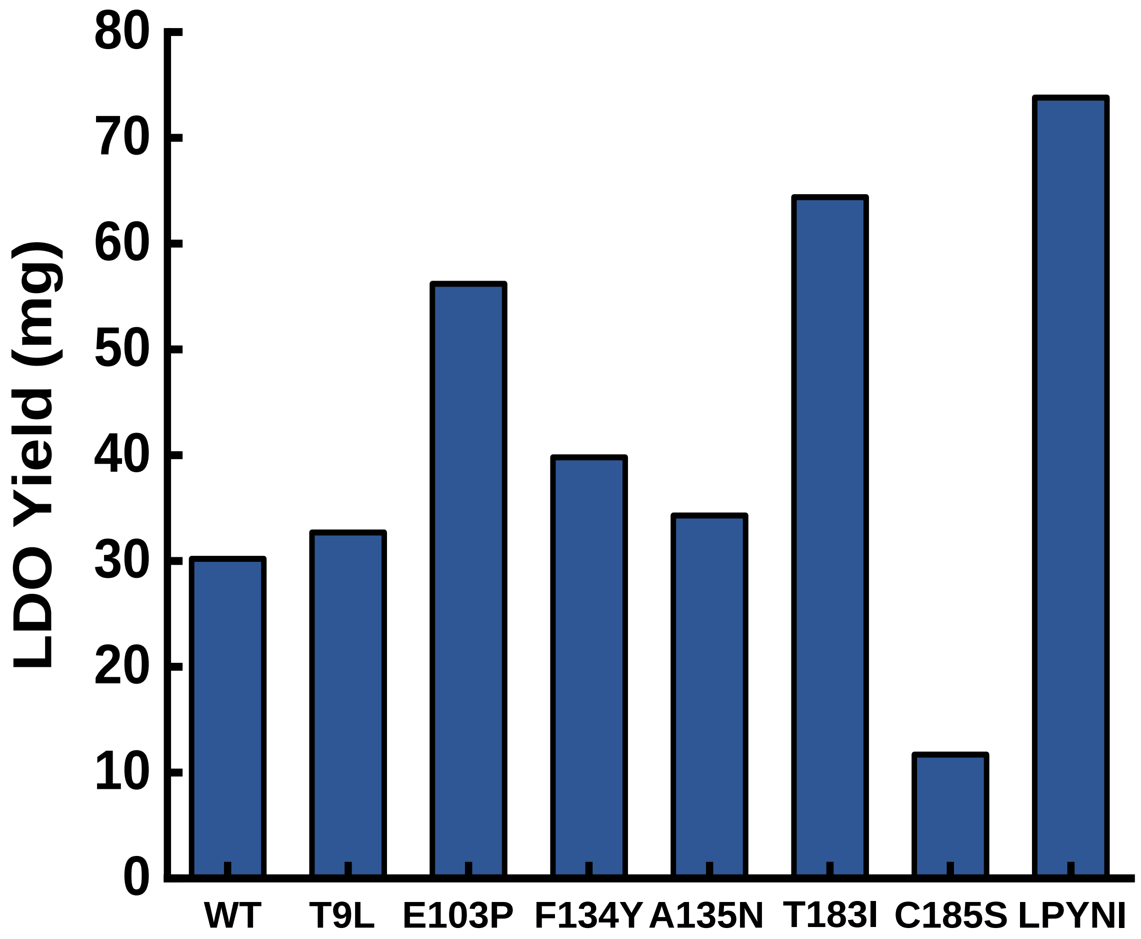


**Figure S6**. Yields of purified, recombinant LDO protein (mg) obtained from 2 L of expressed, cell culture growth. Mutations appear to increase the solubility of the expressed protein, except for C185S.

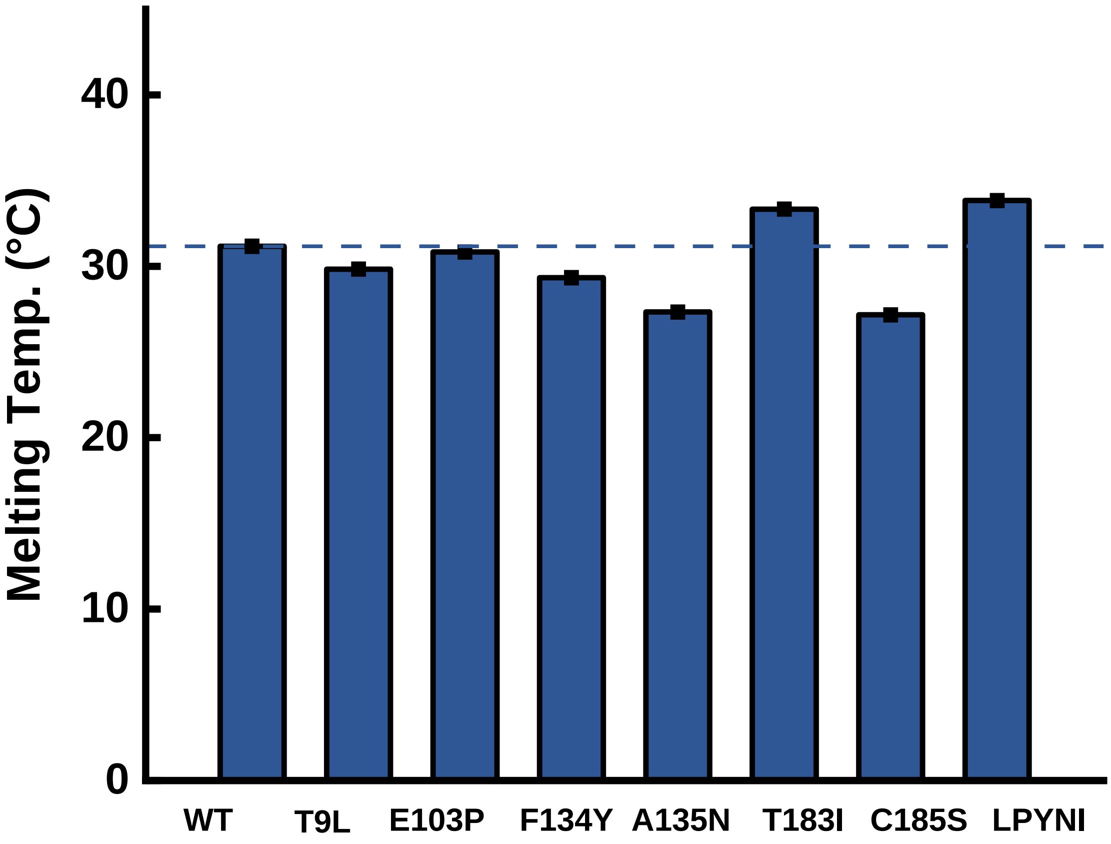


**Figure S7**. Melting point analysis of apo LDO proteins from TSA. Dashed, horizontal line indicates the $T_m$ value for WT hydrox. Error bars are the standard deviation of three biological replicates.

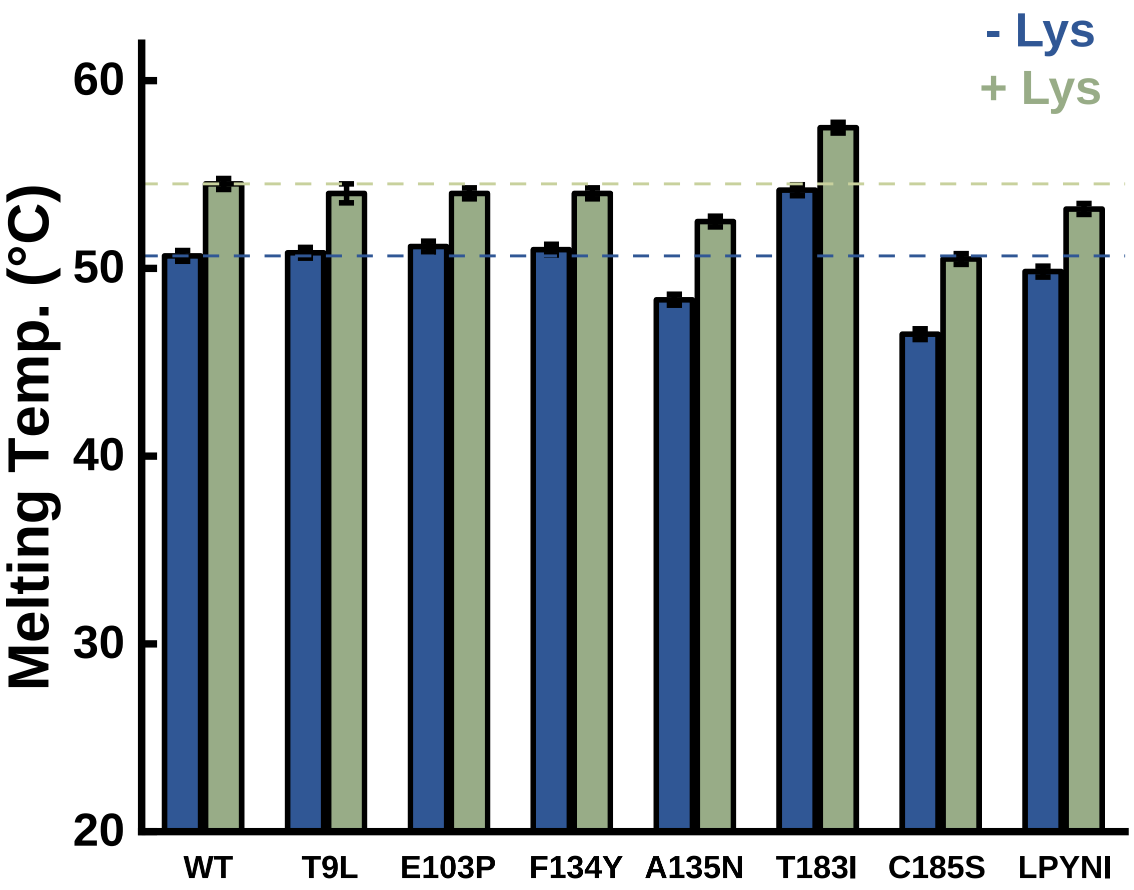


**Figure S8**. TSA melting point analysis of $Mn^{2+}$/2OG-bound LDO proteins without and with Lysine substrate. Dashed, horizontal lines indicates the $T_m$ values for WT Hydrox. Error bars are the standard deviation of three biological replicates.

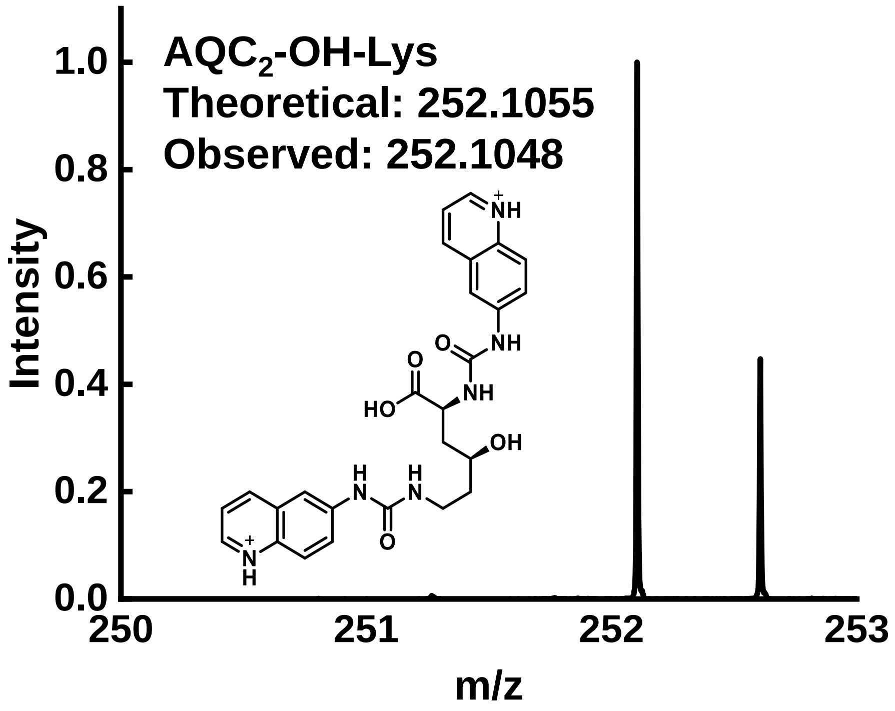


**Figure S9**. Accurate mass measurements for LDO reaction product. Accurate mass spectrum for AQC-tagged OH-Lysine; theoretical $[M+2H]^{2+}$ = 252.1055; mass error = 2.8 ppm.

**Table S3**. Primers used in this work for Site-Directed Mutagenesis. All primers are in 5' → 3' order. Note: All primers were ordered with the 5' nucleotide phosphorylated for compatibility with the Phusion polymerase kit.

| LDO mutant | Forward primer | Reverse Primer |
|---|---|---|
| T9L | GAAATTGACGAGCT GTTGGAGAAGTTC | GTGGACATCCATATG CGGACCTTGGAAC |
| E103P | ACCTTTCTGGCCGGT ATTACGAGAGAG | CAACAGAACTGGTG AACGGGTC |
| F134Y | CCACGGCTGGCATT GGGATGAC | GTGTCAGAGGCATA CTCCTGATGC |
| A135N | CGGCTGGCATTGGG ATGACTACAG | TGGGTGTCAGAGTT GAACTCCTGATG |
| T183I | AGATTGACACCTATG GTCTGGTCTCCG | GACGCTCGCAGAGA ATTTCATTAATACGC |
| C185S | TTGACACCTATGGTC TGGTCTCCG | TCTGACGCTCGCTGA GGGTTTCA |
